## Supplementary material for "Post-mating parental behavior trajectories differ across four species of deer mice": S1 File

### Mating monitoring with the Raspberry Pi Zero

This tutorial describes how to construct a home-made set up to record up to 9 hours of video of a standard Allentown mouse or rat cage (Allentown, Allentown, NJ, USA) in the dark.

#### Material

If provided, follow the link to find the website from which we purchased the item:

1. For the Raspberry Pi
   - [Raspberry Pi Zero version 1.3](https://www.adafruit.com/products/2885)
   - MicroUSB charger (e.g. smartphone charger)
   - [Mini HDMI to HDMI/VGA/DVI cable](https://www.amazon.com/Cable-Rankie-Rated-Speed-Bi-Directional/dp/B00YOSA85Q/ref=sr_1_1?s=electronics&ie=UTF8&qid=1479934402&sr=1-1-spons&keywords=hdmi+to+dvi&psc=1)
   - [USB OTG cable](https://www.adafruit.com/products/1099)
   - [Pi Zero protector](https://www.adafruit.com/products/2883)
   - MicroSD card 16 or 32GB
   - MicroSD card reader
   - Yoobao YB-M4 battery pack or equivalent
   - [2x20 Male header strip](https://www.adafruit.com/products/2822)
   - [USB mini hub](https://www.adafruit.com/products/2991)
   - IR LED strips (3 LEDs per Pi Zero)
2. For the camera
   - [Raspberry Pi Zero v1.3 Camera Cable](https://www.adafruit.com/products/3157)
   - [Pi NoIR camera module](https://www.amazon.com/Raspberry-Pi-NoIR-Camera-Module/dp/B01ER2SMHY/ref=sr_1_1?ie=UTF8&qid=1484532155&sr=8-1&keywords=pi+noir+camera)
   - [18-8 stainless steel hex nuts (thread size: 2-56)](https://www.mcmaster.com/#hex-nuts/=155inaa)
   - [18-8 stainless steel socket head Screws (thread size: 4-40, thread length: 1 3/4")](https://www.mcmaster.com/#standard-socket-head-screws/=14w67lp)
   - Stainless steel socket head screws (thread size: 2-56, thread length does not matter).
   - Fisheye lens for smartphone
   - [1/4"-thick cast acrylic](https://www.mcmaster.com/#standard-plastic-sheets/=17d3u5n)
3. For the Real Time Clock
   - [DS1307 Real Time Clock Assembled breakout board](https://www.adafruit.com/products/3296) and [battery](https://www.adafruit.com/products/380)
   - [1x20 or 2x20 GPIO female header](https://www.adafruit.com/products/2222)
   - Wifi USB dongle

#### Procedure

##### Configuring the SD card

Download the zip file containing the operating system called NOOBS from the [Raspberry Pi website](https://www.raspberrypi.org/downloads/%20NOOBS/) on your computer. Unzip all files. Transfer all files onto the SD card using the micro SD card reader. After this is done, insert the SD card carefully into the Raspberry Pi. Turn on a monitor, plug the Raspberry Pi to the monitor. Plug in a keyboard and a mouse into the USB hub, and the hub into the Pi. Then and only then, plug in the power chord into the Pi. It will turn on automatically.

Follow the instructions on the screen to install the OS image Raspbian Jessie.

##### The Real Time Clock (RTC)

First, solder the male header strip to the Pi zero (S1 Fig); let the board cool down after soldering a few pins, or it may overheat. Then, cut four flexible electric wires (different colours make it easier for the next steps), strip and tin their ends (S2A Fig). Solder one cable to each of the following holes on the RTC assembled breakout board (S2B and C Figs): GND (black), 5V (red), SDA (blue) and SCL (yellow); leave

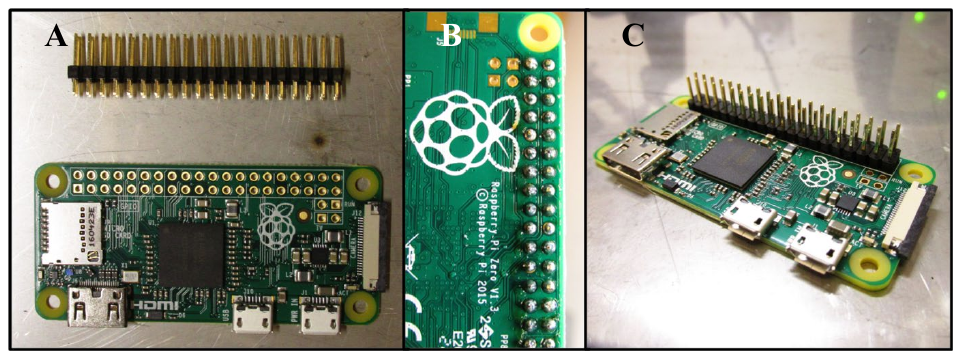

**S1 Fig. Raspberry Pi Zero and soldering of the male header strip. (A)** Raspberry Pi Zero from above and male header strip before soldering. **(B)** Raspberry Pi Zero from below with soldered male header strip. **(C)** Raspberry Pi Zero with soldered male header strip.

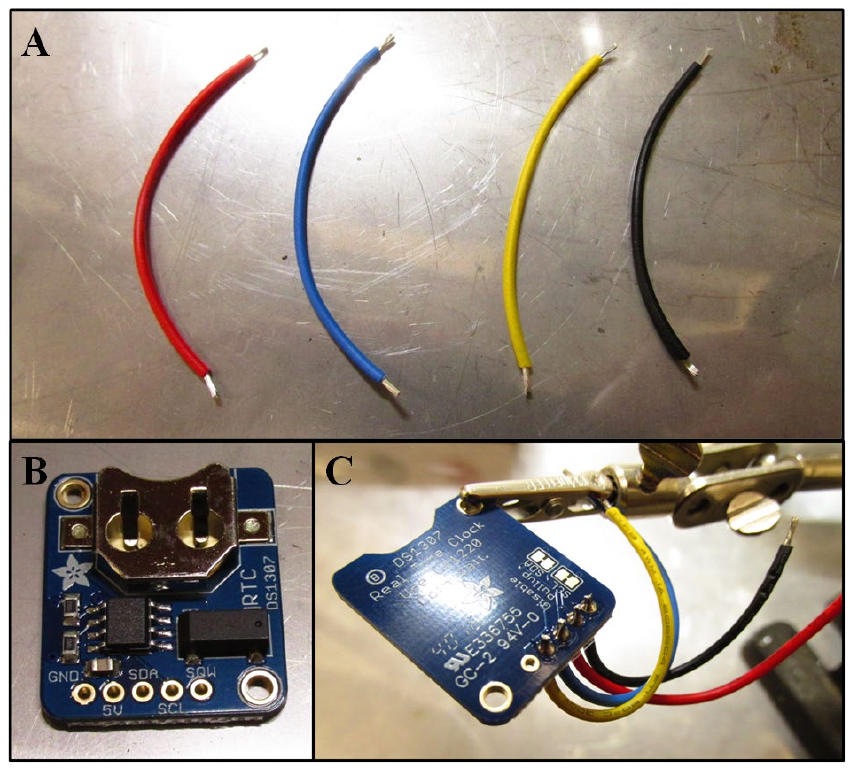

**S2 Fig. Flexible electric wires of different colours and soldering to the Real Time Clock (RTC) assembled breakout board. (A)** The ends of the electric wires are stripped and tinned. **(B)** RTC assembled breakout board. **(C)** Soldering of each colour electric wire to the corresponding hole on the RTC assembled breakout board.

With pliers, cut a 1x2 and a 1x3 section (or one 2x3) of the GPIO female header (S3A Fig), and tin the pins (S3B Fig). Solder the GND and 5V wires to the 1x3, leaving the middle pin untouched (S3C Fig). Solder the SDA and SCL wires to the 1x2 section (S3D Fig). Then, insert the pins of the Raspberry Pi into the GPIO female header (S3E Fig). Also see the distribution of the Pi Zero pins on S4 Fig. The RTC board can then be attached to the Pi's protector with Velcro.

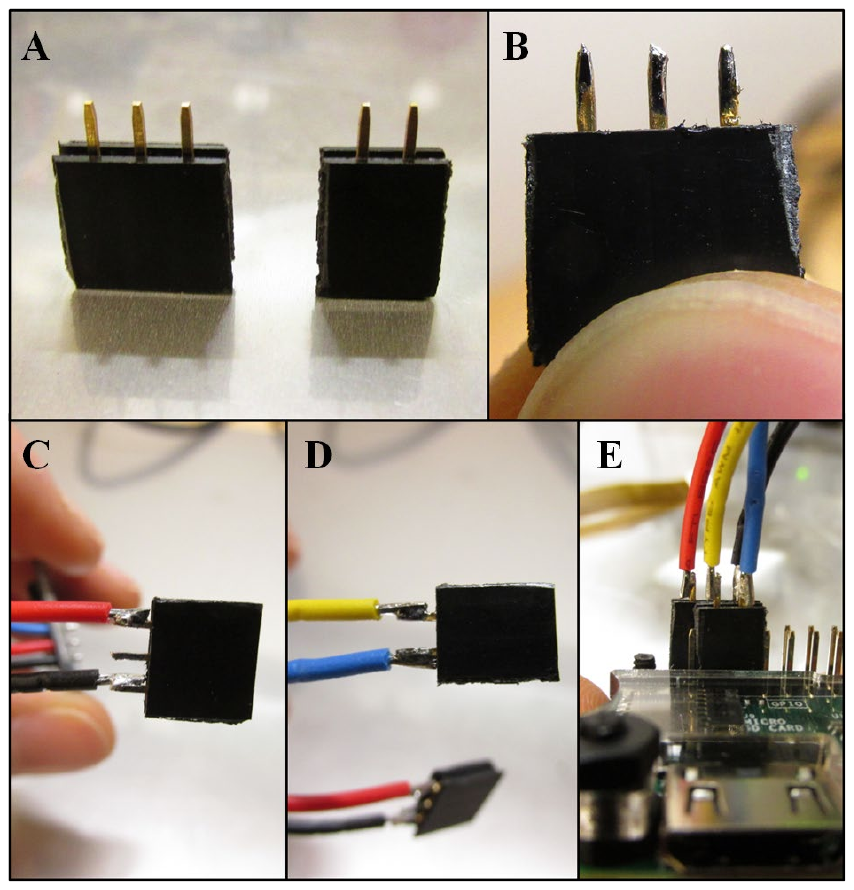

**S3 Fig. GPIO female header and soldering to the electric wires. (A)** One 1x3 and one 1x2 section of a GPIO female header. **(B)** Tinned pins of the 1x3 section. **(C)** Soldering of the 1x3 section to the red wire (5V) and to the black wire (GND). **(D)** Soldering of the 1x2 section to the yellow wire (SCL) and to the blue wire (SDA). Assembly of the electric wires and the GPIO female header to the Raspberry Pi Zero pins.

To configure the RTC, follow the instructions provided on the Adafruit tutorials [here](https://learn.adafruit.com/adafruits-raspberry-pi-lesson-4-gpio-setup/configuring-i2c) (https://learn.adafruit.com/adafruits-raspberry-pi-lesson-4-gpio-setup/configuring-i2c) and then [here](https://learn.adafruit.com/adding-a-real-time-clock-to-raspberry-pi/set-rtc-time) (https://learn.adafruit.com/adding-a-real-time-clock-to-raspberry-pi/set-rtc-time). The wifi dongle must be connected to the Pi in order to set the time precisely. While configuring the RTC, its battery should be inserted. The time zone of the RTC can be changed by opening a terminal, typing in sudo raspi-config, and then follow Internationalization.

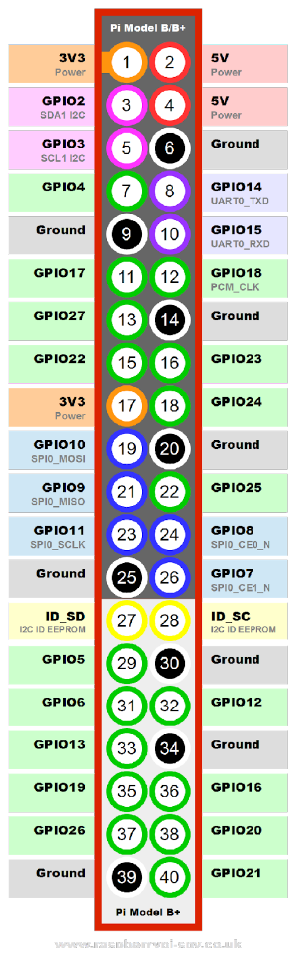

**S4 Fig. Pin distribution of the Raspberry Pi models 3B and Zero from http://www.jameco.com/Jameco/workshop/circuitnotes/raspberry-pi-circuit-note.html.** The pins that are of use in this tutorial are 5V, Ground (GND), GPIO2 SDA I2C (SDA) and GPIO3 SCL1 I2C (SCL).

##### The camera

The camera will come with a camera cable that is only suitable for larger models of the Raspberry Pi. Thus, it needs to be unplugged. To do so, gently pull the black piece of plastic that is maintaining the cable connected to the camera. Pull on both sides simultaneously and be careful not to pull it off completely. The cable should slide out of the slit. Replace it with a cable compatible with the Pi Zero v1.3. Connect the cable to the Pi Zero while the latter is off. When this is done, turn on the Pi and in a terminal, type in sudo raspi-config and follow Enable the camera.

##### Camera holder

The camera holder has to be laser-cut in a piece of 1/4"-thick cast acrylic. The holders for both Allentown mouse and rat cages were designed with Autodesk Inventor Professional 2016 (<http://www.autodesk.com>). The drawings are provided as separate files ("CameraHolder_MouseCage.dxf", "CameraHolder_RatCage.dxf"). In the cut-out piece of acrylic, insert the fisheye lens. Depending on the model, a file will have to be used to slightly increase the size of the hole. Test it regularly after a bit of filing, until the lens fits.

For the camera to be attached to this piece, two 2-56 screws need to be used. Do not screw them all the way in, leave ~2mm, otherwise the camera will be tilted. Secure the screws with two hex nuts.

Use a 1-18 screw in the lower hole of the holder, it will later be useful to adjust the angle of the camera relative to the cage wall.

##### LED strip

To provide enough IR light to the camera, you need to prepare an LED strip. Three LEDs are enough for mouse and a rat cages, but you need to jump all the resistors by soldering a small piece of wire onto them (S5A Fig). The strip also needs a connector cable. To connect the LED strip to the Pi, it is necessary to solder the connector cable to a USB cable. To do so, strip the USB cable, cut the green and white wires short, but at different lengths to avoid contact. Strip the red and black wires. Slide one large heat-shrinkable tube around the whole USB cable. Strip the connector cable (both red and black parts). Slide each wire into a smaller heat-shrinkable tube. Tin and solder the ends of the connector cable to the ends of the USB cable. After this is done, heat-shrink the small tubes, and then the larger tube over them, to protect the connection (S5B Fig).

You will then be able to plug in the LED strip to the USB OTG cable (S5C Fig), and to the Pi zero. Check that the LEDs work. They should provide enough light for a mouse or a rat cage.

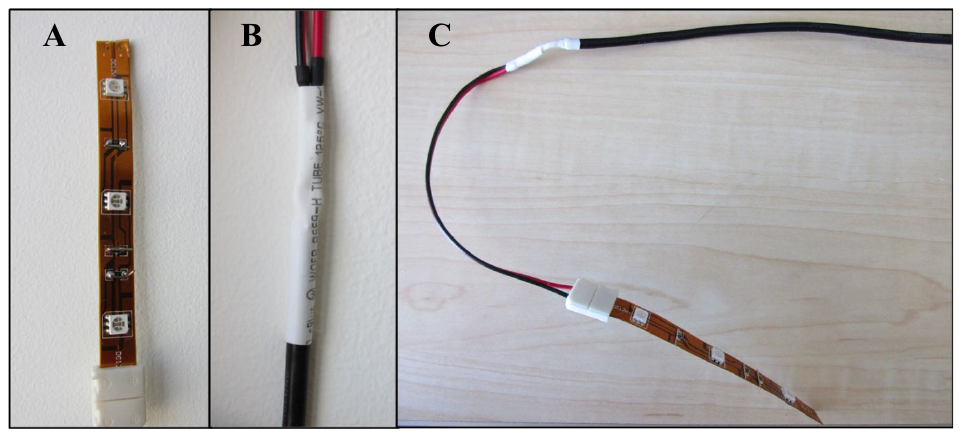
**S5 Fig. Modification of an IR LED strip to illuminate the cage. (A)** LED strip with jumped resistors. **(B)** Heat-shrinkable tubes connecting the USB cable to the connector cable. **(C)** Final assembly of the LED strip and USB cable.

##### Assembly

The Pi zero should be on top of the cage's lid. The camera cable should come towards the cage card holder, be slid into the slit between the green rectangle and the metallic piece of the card holder (S6A Fig), and then connected to the camera (S6B and C Figs). This is a delicate procedure. The camera setup should be attached to the card holder. Then you can adjust the angle of the camera with the long 1-18 screw (S6D and E).

To start up the Pi, simply plug in the battery pack.

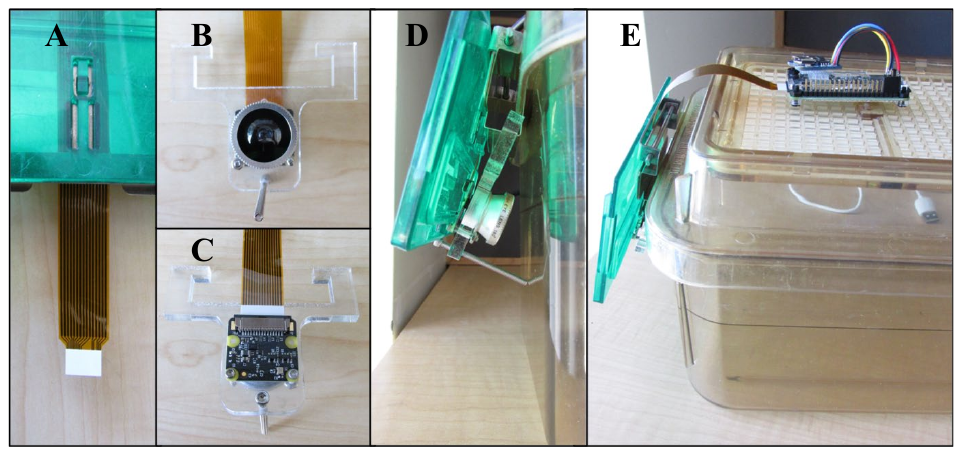
**S6 Fig. Final assembly of the camera set up on a mouse cage. (A)** The camera cable goes into the slit of the cage card holder. **(B)** Assembled camera, fisheye lens, camera cable, and screw (front view). **(C)** camera, fisheye lens, camera cable, and screw (back view). **(D)** Final assembly on a mouse cage, the long screw allows to adjust the angle of the camera (side view). **(E)** Final assembly on a mouse cage, the Raspberry Pi Zero is on top of the cage, the camera cable is connected to the Pi and to the camera. The battery pack is not on the picture.

##### Programming the Pi to record at a given time

First, create a folder on the desktop with an informative name ("Mating_monitoring" in our case). Then, on the desktop, paste the python script provided below. This will record 8hrs 15mins of videos in 5-minute chunks and store them in a .h264 format in the Mating_monitoring folder. You can read this format with VLC (<http://www.videolan.org/vlc>), or you can convert them to .mp4 with these extra lines to the python script:

conversion = "MP4Box -fps 30 -add xxxx.h264 xxxx.mp4"

call ([conversion], shell=True)

deletion = "rm xxxx.h264" # optional if you want to delete the .h264 video and keep only the converted file

call ([deletion], shell=True)

Then, open a terminal, and type in:

crontab -e

This will open a file that allows to program the launching of scripts at given time points. Instructions are given in the file. At the bottom of the file, type in, for example:

55 13 * * * python home/pi/Desktop/Mating_monitoring.py

15 22 * * * sudo shutdown -h now

The first line launches the python script that will record videos at 13:55. The second line allows the Pi to shutdown once the recording is done, at 22:15. It is better for the lifespan of the SD card to shut down by a command than by running out of battery.

Script:

```
from picamera import PiCamera
from time import sleep
from time import strftime

# Configuration
resolution = (640, 480)
hours_of_video = 8.25
segment_length = 5 # in minutes, allows to record videos in 5-minute chunks

# Run Camera
camera = PiCamera(resolution = resolution)
camera.rotation = 180

# camera.start_preview() # if you want to visualize recording as it goes

# Number of segments
segments = int(hours_of_video*60/segment_length)

for i in range (segments):
    date_string = strftime('%Y-%m-%d_%H-%M')
    camera.start_recording('/home/pi/species_pairID_' + date_string + '.h264') # numbers videos automatically
    camera.wait_recording (segment_length*60) # in seconds
    camera.stop_recording()
# camera.stop_preview()
```

##### Recording and storing videos

Once the python script, the desktop folder, and the crontab are ready, you will just have to plug in the Pi once it is set up on a cage. The recording will start as specified in crontab.

Once the recording is done, you can unplug the Pi, take out the SD card, slide it in a Raspberry Pi 3 (faster than Pi Zero when transferring many files) previously connected to a monitor. Plug in the Raspberry Pi 3 and transfer the videos from the Mating_monitoring folder onto a USB stick or external hard drive.
