## Supplementary figures and images for "Post-mating parental behavior trajectories differ across four species of deer mice"

### S7 Fig

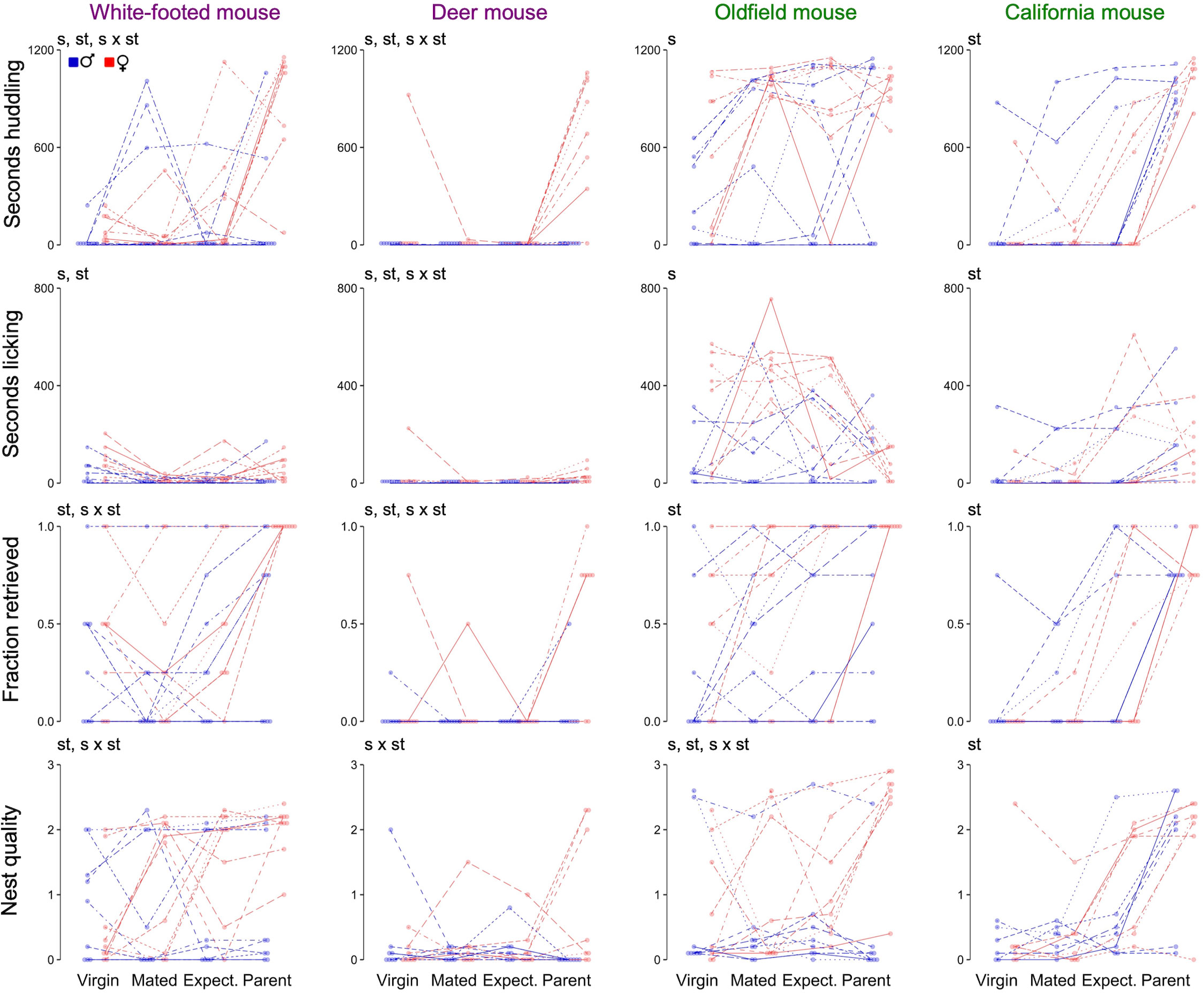
